# Optimized Enrichment of Murine Blood-Brain Barrier Vessels with a Critical Focus on Network Hierarchy in Post-Collection Analysis

**DOI:** 10.1101/2024.09.19.613898

**Authors:** Hanaa Abdelazim, Audra Barnes, James Stupin, Ranah Hasson, Carmen Muñoz-Ballester, Kenneth L. Young, Stefanie Robel, James W. Smyth, Samy Lamouille, John C. Chappell

## Abstract

Cerebrovascular networks contain a unique region of interconnected capillaries known as the blood-brain barrier (BBB). Positioned between upstream arteries and downstream veins, these microvessels have unique structural features, such as the absence of vascular smooth muscle cells (vSMCs) and a relatively thin basement membrane, to facilitate highly efficient yet selective exchange between the circulation and the brain interstitium. This vital role in neurological health and function has garnered significant attention from the scientific community and inspired methodology for enriching BBB capillaries. Extensive characterization of the isolates from such protocols is essential for framing the results of follow-on experiments and analyses, providing the most accurate interpretation and assignment of BBB properties. Seeking to aid in these efforts, here we visually screened output samples using fluorescent labels and found considerable reduction of non-vascular cells following density gradient centrifugation (DGC) and subsequent filtration. Comparatively, this protocol enriched brain capillaries, though larger diameter vessels associated with vSMCs could not be fully excluded. Protein analysis further underscored the enrichment of vascular markers following DGC, with filtration preserving BBB-associated markers and reducing – though not fully removing – arterial/venous contributions. Transcriptional profiling followed similar trends of DGC plus filtration generating isolates with less non-vascular and non- capillary material included. Considering vascular network hierarchy inspired a more comprehensive assessment of the material yielded from brain microvasculature isolation protocols. This approach is important for providing an accurate representation of the cerebrovascular segments being used for data collection and assigning BBB properties specifically to capillaries relative to other regions of the brain vasculature.

**HIGHLIGHTS:**

– We optimized a protocol for the enrichment of murine capillaries using density gradient centrifugation and follow-on filtration.
– We offer an approach to analyzing post-collection cerebrovascular fragments and cells with respect to vascular network hierarchy.
– Assessing arterial and venous markers alongside those associated with the BBB provides a more comprehensive view of material collected.
– Enhanced insight into isolate composition is critical for a more accurate view of BBB biology relative to larger diameter cerebrovasculature.

**MOTIVATION:** The recent surge in studies investigating the cerebrovasculature, and the blood-brain barrier (BBB) in particular, has inspired a broad range of approaches to target and observe these specialized blood vessels within murine models. To capture transcriptional and molecular changes during a specific intervention or disease model, techniques have been developed to isolate brain capillary networks and collect their cellular constituents for downstream analysis. Here, we sought to highlight the benefits and cautions of isolating and enriching microvessels from murine brain tissue. Specifically, through rigorous assessment of the output material following application of specific protocols, we presented the benefits of specific approaches to reducing the inclusion of non-vascular cells and non-capillary vessel segments, verified by analysis of vascular-related proteins and transcripts. We also emphasized the levels of larger-caliber vessels (i.e. arteries/arterioles and veins/venules) that are collected alongside cerebral capillaries with each method. Distinguishing these vascular regions with greater precision is critical for attributing specific characteristics exclusively to the BBB where metabolic, ion, and waste exchange occurs. While the addition of larger vessels to molecular / transcriptional analyses or follow-on experiments may not be substantial for a given protocol, it is essential to gauge and report their level of inclusion, as their contributions may be inadvertently assigned to the BBB. Therefore, we present this optimized brain microvessel isolation protocol and associated evaluation methods to underscore the need for increased rigor in characterizing vascular regions that are collected and analyzed within a given study.

**Graphical Abstract:** Constructed using resources from BioRender.com.

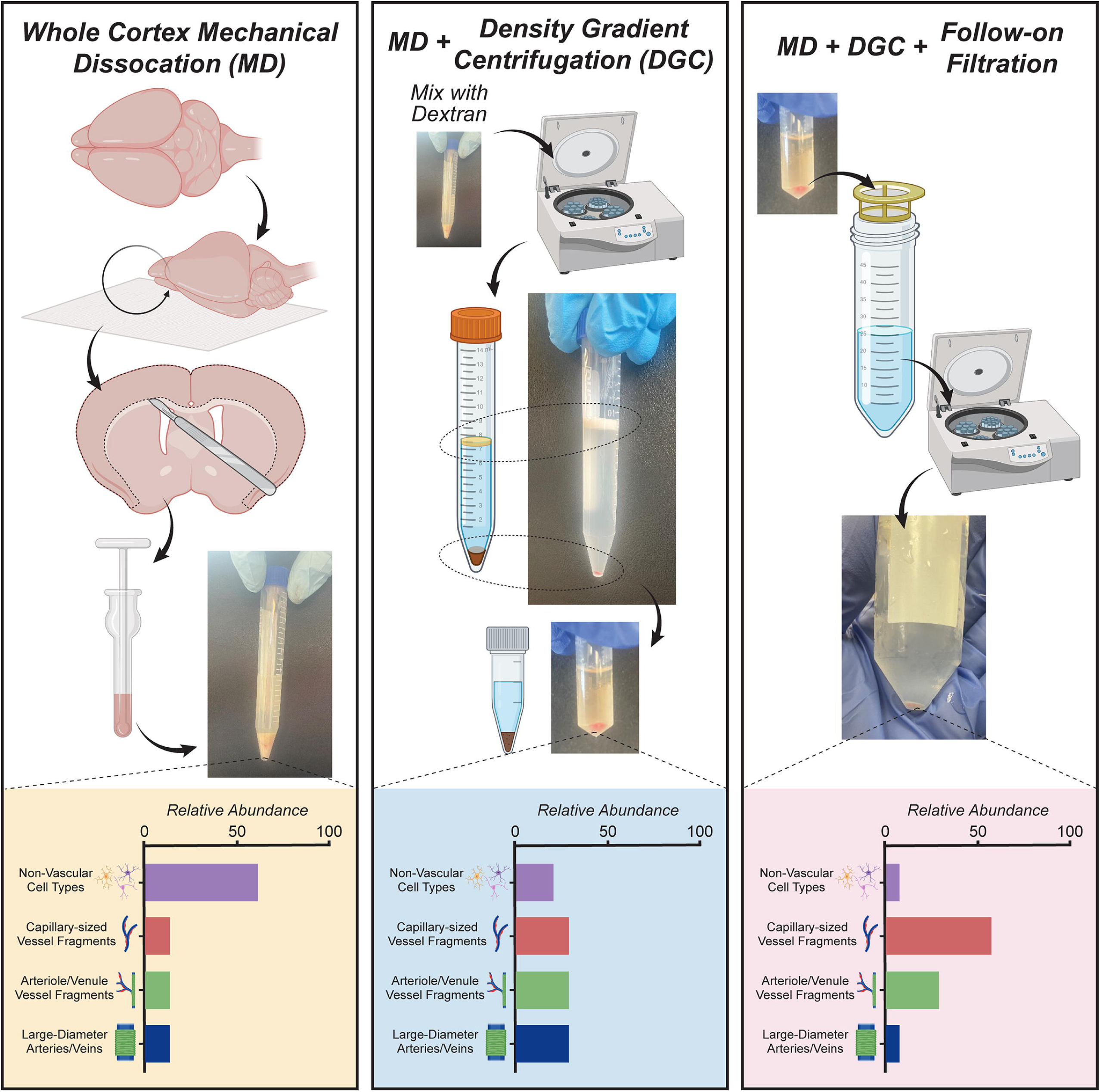

## INTRODUCTION

The blood vasculature is a continuum of interconnected conduits, distributing nutrients, hormones, and cells systemically while also removing waste and metabolic byproducts, among other critical roles. To accomplish these functions, the vascular system contains regions that share key similarities but also important distinctions on the cellular, molecular, and morphological levels^1^. Arteries and arterioles, for example, are generally composed of: (i) an inner lining of endothelial cells (ECs), (ii) a relatively dense extracellular matrix (ECM), or basement membrane, to withstand intraluminal pressures, and (iii) corresponding layers of concentrically wrapped vascular smooth muscle cells (vSMCs) that dynamically regulate vessel diameter and tissue perfusion^2^. Downstream capillary networks are highly branched and have relatively thin walls to facilitate adequate blood-tissue exchange of metabolites, ions, and waste, among other factors. Depending on the organ or a subregion therein, endothelial permeability can be classified as continuous, fenestrated, or discontinuous, with corresponding levels of abluminal pericyte (PC) association and ECM deposition^3^ Capillaries then feed into venules, which merge into veins. Venous regions contain more loosely associated ECs, cobblestone-like vSMCs, and a unique ECM composition, which collectively facilitate immune cell trafficking but also passive distension for blood volume reserve^4^. Thus, while the vasculature is generally comprised of an endothelium, mural cells (vSMCs/PCs), and an ECM basement membrane, these constituents exhibit highly specific characteristics that align with their region- and tissue-specific function, as highlighted by recent insights into EC arteriovenous specification^1,5–7^ and vascular heterogeneity across different tissues and organs^8,9^.

With continually increasing interest in neuroscience research, brain blood vessels have also garnered significant attention of late, though their importance has been recognized for some time^10^. Interest has flourished in utilizing the cerebrovasculature to treat neurological conditions and brain-specific cancers as well as in better understanding pathologies that directly involve cerebrovascular dysfunction such as stroke and Alzheimer’s Disease^11^ This recent focus on the brain vasculature has revealed distinct properties within a subset of these vessels denoted as the blood-brain barrier (BBB).

This semi-permeable barrier is highly selective in regulating which solutes can traverse the capillary wall into and out of the brain parenchyma. Because capillaries are the primary site for metabolic, ion, and waste exchange, most studies attribute BBB characteristics exclusively to the cerebral capillaries and particularly those not found within circumventricular organs, the choroid plexus, or dura mater^12^. This specialized barrier function is suggested to arise from: (i) a unique augmentation of EC tight junctions (e.g. Claudin5/*Cldn5*-enriched), (ii) PC incorporation within microvessel walls, (iii) a thin deposition of specific ECM components, and (iv) astrocyte end-foot apposition along the abluminal surface. While the concept has recently been suggested^13^, arterioles and venules are generally not assigned to the BBB. Metabolic exchange is limited for these vessels owing to the presence of vSMCs and a thicker basement membrane, though cerebral artery and vein endothelium may exhibit BBB-like properties. Therefore, since brain capillary networks are largely considered to represent the BBB, it is critical to develop rigorous protocols to isolate and enrich these cerebral microvessels for downstream analysis. Alongside this goal, methods must also be developed to assess the levels of non-capillary vasculature that may not be completely excluded by a specific isolation technique, as that material will influence the output data and any subsequent interpretations of BBB properties.

Given the extensive metabolic demands of the brain and its wide array of functions and cellular components, the cerebrovasculature is tasked with meeting these demands on a continual basis. It does so despite the modest number of vascular cells relative to the abundance of other cell types comprising the over 80 billion cells within the human brain^14^ Analyzing molecular changes within this cerebrovascular compartment and its various subregions can be challenging to nearly impossible without protocols to enrich associated vascular cell types. Single cell transcriptomics have addressed this challenge to a certain extent, but this approach must also be coupled with methods to validate gene expression data on mRNA, protein, and morphological levels^15–17^. Approaches such as magnetic-activated cell sorting (MACS) or fluorescence-activated cell sorting (FACS) utilize labeling by reporter constructs or cell surface markers that may not be restricted exclusively to brain capillary-associated cells^18,19^. Laser capture microdissection allows cells to be isolated from cerebrovascular segments based on their morphology, but this technique suffers from low yield, limited regional sampling, and significant time investment^20^ Recent protocols have been developed to process mechanically dissociated brain tissue using density gradient centrifugation (DGC), with varying use of filtration techniques, to select capillaries based on size^21–27^ These methods have shown great promise in improving the enrichment of capillaries from brain tissue, but opportunities exist to enhance follow-on assessment of the resulting material, especially in accounting for vascular network hierarchy.

Various strategies have recently emerged for dissociating cerebral tissue into its structural and cellular components, separating the contents based on physical properties, and analyzing the collection output for BBB characteristics and the inclusion or exclusion of larger diameter vessels. Applying filter paper to specific brain surfaces has emerged as a clever approach to remove unwanted structures such as the meninges and/or choroidal plexus vessels^22,25,26^. Moreover, depending on the area of interest, surgical dissection of the cortex has also proven to be relatively easy and efficient for enriching vessels from this region. Mechanical dissociation via mortar-pestle or Dounce grinding is often preferred over serial needle shearing, which may risk damage on the molecular level. Follow-on enzymatic digestion via collagenase or trypsin has been explored for releasing individual cells^22,24^, though concerns with generalized cell activation exist with this approach. Separating disaggregated brain tissue into distinct components can be achieved in part based on unique physical properties, especially when mixed with certain reagents. Dextran, Ficoll, and bovine serum albumin (BSA) have all been explored for generating distinct layers during centrifugation^21–27^, leading to a gradient of highly dense, lipid-rich structures such as myelin aggregating at the top and predominantly vascular structures settling to the bottom. Follow-on filtration of this vascular portion can further enrich -- *though not fully purify --* microvessels that largely represent the BBB.

Because these cerebrovascular fractions will inevitably contain some level of non-capillary vessels, it is critical to rigorously characterize the collection output and frame data interpretation with this insight. Standard approaches often include assessment of BBB-associated morphology and markers alongside profiling of non-vascular cells^23–25^; interestingly, some EC junction molecules suggested to define the BBB (e.g. *Cldn5, Ocln, Pecam1, Cdh5*) are also detected at relatively high levels within the endothelium of cerebral arteries/arterioles and veins/venules^28–30^. Furthermore, morphological analysis may also extend towards the use of SMC markers such as α-smooth muscle actin (αSMA, gene: *Acta2*)^21,26^ but a more comprehensive evaluation of larger vessels, particularly by transcriptional profiling (e.g. *Myh11*, *EphB4*), is often limited or absent. Here, we drew from these emerging approaches and optimized a protocol to enrich cerebral capillaries that are largely considered representative of the BBB. In addition, we applied a suite of analyses to account for vascular network hierarchy and gauge how distinct steps in this protocol affected inclusion of non-vascular cells and larger diameter vessels in the output material.

## RESULTS

### Visualizing Cerebrovascular Fragments from Different Isolation Protocols Reveals Heterogeneity in Vessel Caliber and Non-vascular Cells in Collection Outputs

Recent interest in studying the blood-brain barrier (BBB) has yielded a broad range of methods to collect cerebral microvessels. Varying approaches have been applied to characterize the collection output, specifically regarding inclusion of non-capillary sized vessels and non-vascular cells. At the outset of our own BBB-focused study, we sought to compare different protocols for isolating and enriching brain capillaries (Figure 1), knowing that few, if any, higher-yield strategies are capable of fully excluding larger caliber vessels like arterioles and venules. We reasoned that it would be important to have a clear understanding of how much non-capillary structures might contribute to our downstream data analysis and to any interpretations of BBB properties. To begin this comparison, we utilized animals expressing a fluorescent reporter (DsRed) driven by the promoter of a well-accepted vascular mural cell gene, *Cspg4*/NG2 (expressed in PCs and vSMCs). We co-labeled endothelial cells (ECs) with isolectinB4 conjugated to AlexaFluor647. Follow-on staining with DAPI also facilitated visualization of nuclei within vascular and non-vascular cell types. High-resolution confocal imaging revealed non-vascular cells within tissue fragments derived from whole brain input that was mechanically dissociated by mortar-pestle grinding and DGC, but without filtration (Figure 2A). We then visually assessed microvascular fragments that were collected by DGC and 100μm-pore filtration (see Figure 1). These vessels were antibody labeled against (i) platelet-endothelial cell adhesion molecule-1 (PECAM-1/CD31) for ECs, (ii) platelet-derived growth factor receptor-β (PDGFRβ/CD140b) for vascular mural cells i.e. SMCs and pericytes (PCs), and (iii) αSMA, which is found at high levels predominantly in SMCs. DAPI labeling facilitated identification of vascular and non-vascular cell nuclei (Figure 2B). Confocal images of these samples showed a notable reduction in non-vascular cells. This microvessel enrichment procedure also yielded distinct PC-associated capillary fragments alongside a modest population of αSMA+ vessels with slightly larger diameters. These visualization strategies also confirmed that these protocols could conserve anatomical structures with even the most complex methodology, allowing assessment of which type/size of vessels were being collected within each fraction.

**Figure 1.**
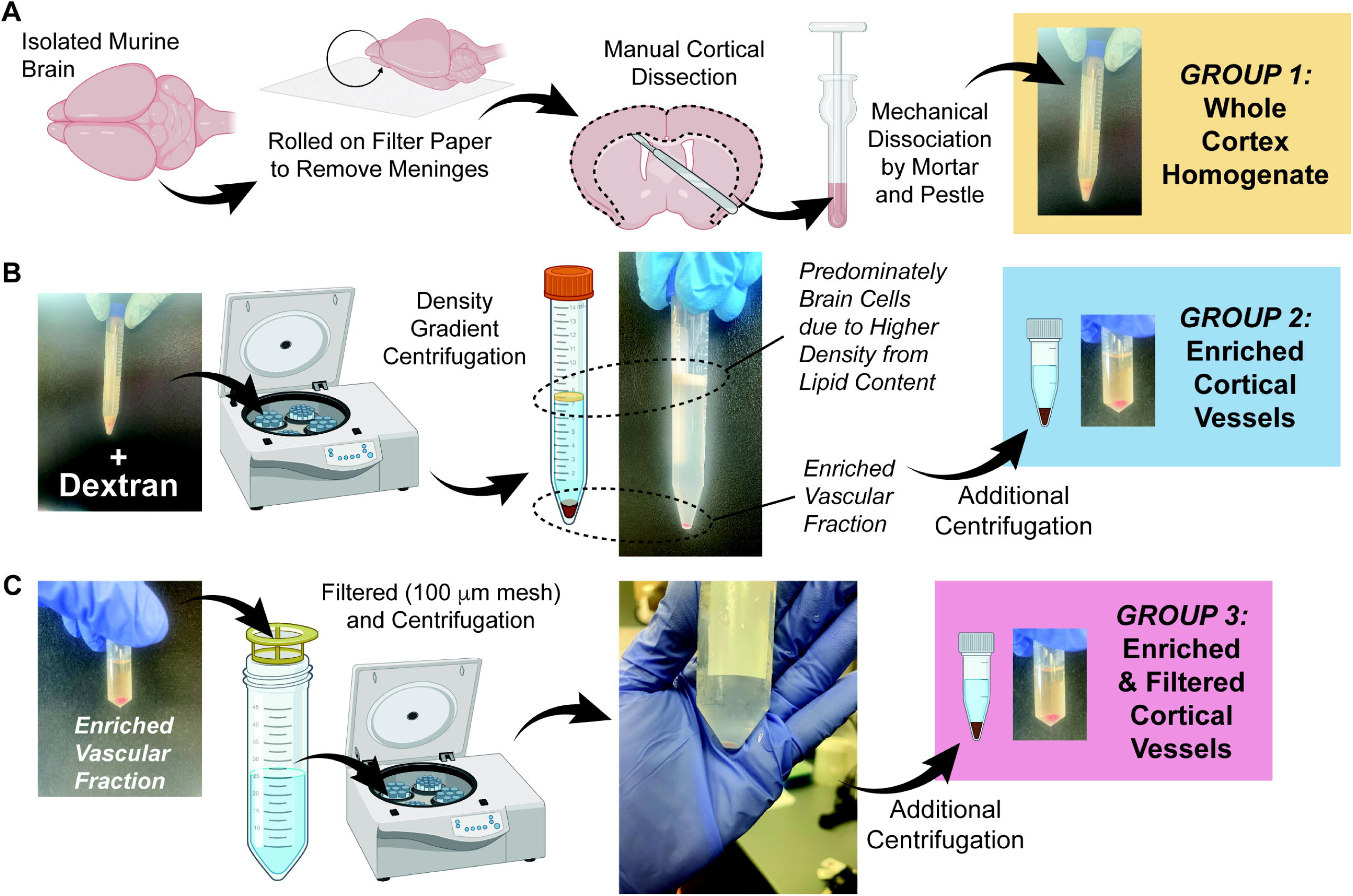
Schematic overview of the enrichment of blood-brain barrier vessels from murine cortex. (A) Schematic of protocol steps applied to remove meninges from isolated murine brains, isolate cortical regions through manual dissection, and mortar- pestle-mediated mechanical dissociation, yielding the whole cortex homogenate representing Group 1. Representative image of resultant material in a 15 mL conical tube is shown. (B) Schematic of additional protocol steps used to process whole cortex homogenate and enrich cortical vasculature by mixing with Dextran and establishing a density gradient by centrifugation, generating a vascular fraction. An additional centrifugation step facilitated final isolation of enriched cortical vessels (Group 2), shown in the representative image. (C) Schematic of follow-on filtration and centrifugation of enriched cortical vessels to produce Group 3 containing enriched and filtered cortical microvessels. Figure constructed using resources from BioRender.com.

**Figure 2.**
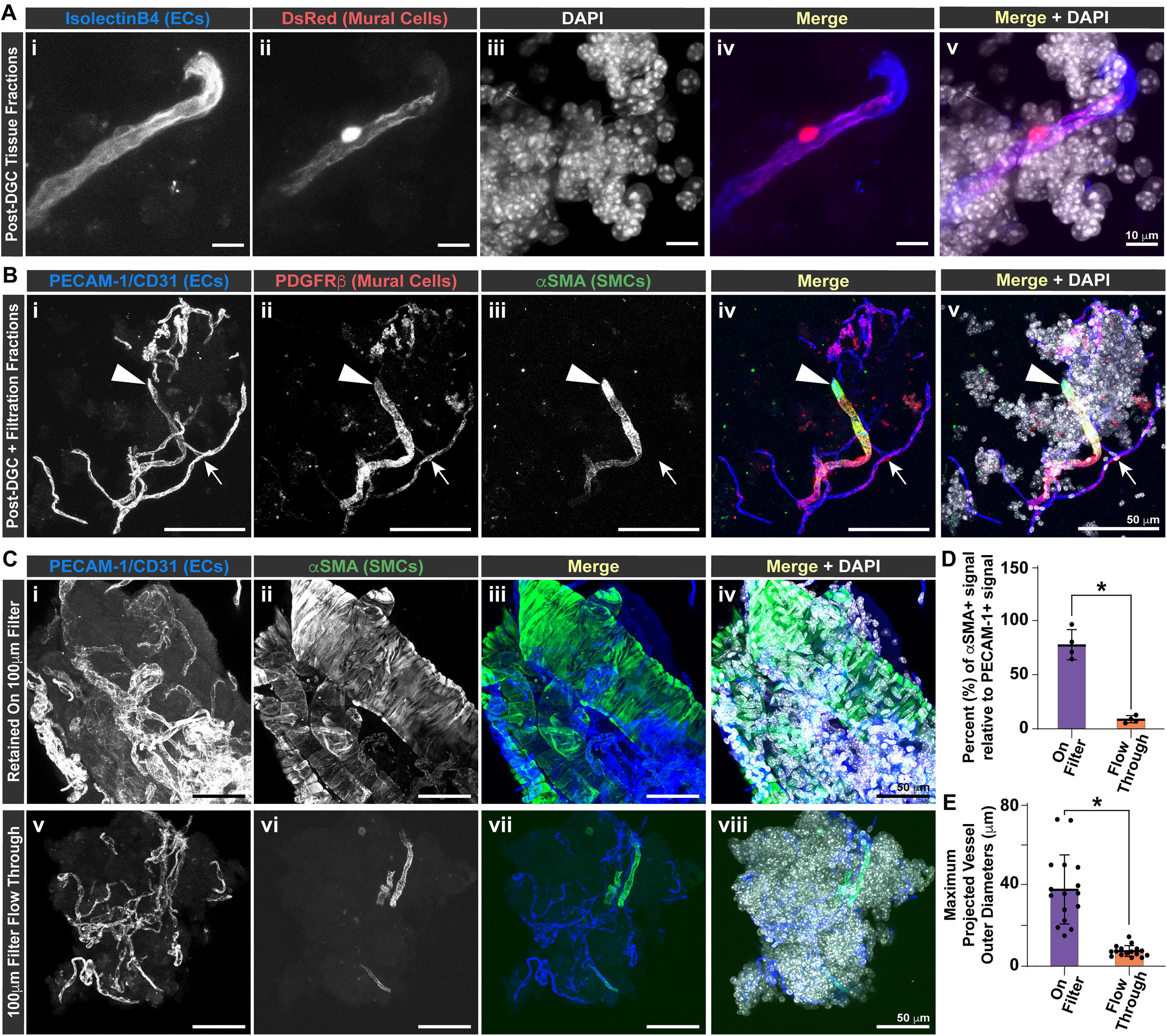
High-power Confocal Imaging of Isolated Brain Microvessels Allows Visual Assessment of Vessel Size and Composition Along with Density of Non- vascular Cells Following Specific Protocol Steps. (A) Representative confocal images of adult mouse brain tissue fractions following DGC without filtration, labeled by IsolectinB4 (i, blue in iv and v), and DsRed from the *Cspg4*/NG2 promoter (ii, red in iv and v), with cell nuclei stained by DAPI (white in iii and v). Scale bars are 10 microns. (B) Representative confocal images of adult mouse brain tissue fractions following DGC and filtration labeled for PECAM-1/CD31 (i, blue in iv and v), PDGFRβ/CD140 (ii, red in iv and v), and αSMA (iii, green in iv and v) with cell nuclei stained by DAPI (white in v). Scale bars are 50 microns. The arrowhead denotes a larger diameter vessel positive for all 3 vascular markers, while the arrow points to a smaller diameter vessel labeled only by PECAM-1 and PDGFRβ, highlighting the mixed population of vessels after filtration. (C) Representative confocal images of adult mouse brain tissue, a fraction of which was retained on a100-micron filter (i-iv) and another fraction that flowed through this filter (v-viii). Cortical vessels are labeled for PECAM-1/CD31 (i and v, and blue in iii, iv, vii, and viii) and αSMA (ii and vi, and green in iii, iv, vii, and viii) with cell nuclei stained by DAPI (white in iv and viii). Scale bars are 50 microns. Note the presence of αSMA-negative, capillary-sized vessels alongside several αSMA+ larger vessels in the on-filter images (i- iv) compared to the flow-through portion (v-viii) containing mostly αSMA-negative, capillary-sized vessels with relatively few αSMA-positive vessel segments. (D) Graph of the percentage of the projected αSMA+ signal area relative to the projected PECAM-1+ signal area for images taken of vessel fragments as shown in (C). Measurements were taken from the on-filter and flow-through portions. Individual data points represent the averages of technical replicate measurements from each biological replicate (n=4 biological replicates). Averages shown by the purple (on-filter) and orange (flow-through) bars were calculated from the biological replicate averages. Error bars are standard deviation, and * denotes p<0.05 by Student’s T-test. (E) Graph of the maximum projected vessel outer diameter based on PECAM-1-labeling of vessel fragments as shown in (C). Measurements were taken from the on-filter and flow-through portions. Individual data points represent the technical replicate measurements from each biological replicate (n=4 biological replicates, with 4 measurements from each). Averages shown by the purple (on-filter) and orange (flow-through) bars were calculated from the technical replicate measurements. Error bars are standard deviation, and * denotes p≤0.05 by Student’s T-test.

From these initial observations, we hypothesized that the vessels retained on the 100μm-pore filter were mostly larger caliber, containing a higher percentage of αSMA+ cells, potentially of value for studies not focused on the BBB and cerebral microcirculation. To capture the morphological differences between the retained vessels and those within the flow-through, we retrieved each fraction and applied antibodies against PECAM-1/CD31 and αSMA as before (plus DAPI), with subsequent high-resolution confocal imaging (Figure 2C). Assessing these images for the percentage of αSMA+ vessels relative to the total PECAM-1+ signal, we found that the on-filter vessels indeed exhibited a significantly higher level of αSMA+ label compared to the flow- through fraction (Figure 2D). Furthermore, the on-filter vessels displayed a wider range of projected outer diameters, with an average diameter of about 40μm; in contrast, vessels within flow-through samples had a narrower diameter range and an average projected outer diameter of about 10μm (Figure 2E). Taken together, these visualization strategies confirmed that (i) DGC (i.e. enrichment) and filtration reduced the inclusion of non-vascular cells in output samples, and (ii) 100μm-pore filtration improved yields of capillary-sized vessels with some smaller diameter, αSMA+ microvasculature also collected.

### Density Gradient Centrifugation of Mechanically Dissociated Murine Cortex Enriches Vascular-Related Proteins with Follow-on Filtration Reducing Larger Vessel SMC Contributions

To complement the image-based assessment of cortical vessel isolation protocols, we also collected protein lysates from the output material of each method. These lysates were processed for western blot to compare levels of vascular-related proteins, specifically: (1) αSMA, a contractile protein predominantly found in vascular SMCs of larger-caliber arteries/arterioles and veins/venules, (2) Claudin5, a tight junction molecule between ECs within BBB capillaries but also arteries and veins^28^, and (3) PECAM-1, a nearly ubiquitous EC adhesion protein present throughout most vascular networks. We utilized α-tubulin as a housekeeping marker across all three collection protocols. Our western blot images revealed a substantial increase in the concentration of all three vascular-related proteins in samples acquired from DGC enrichment as compared to cortex isolation alone (Figure 3A, see Supplementary Figures 1 and 2 for original blots). These increases were confirmed by subsequent image analysis, that is, calculating the ratios of integrated densities between each target signal and the housekeeping marker for whole cortex and enriched samples (Figure 3B). Lysates generated by enrichment plus filtration contained less SMC-associated αSMA protein relative to the levels yielded by enrichment alone; in contrast, Claudin5 and PECAM-1 densities remained elevated compared to whole cortex (Figure 3). These data suggest that, compared to collecting protein directly from mechanically dissociated murine cortex, vessel isolation protocols involving enrichment and filtration yield a similar enhancement of EC-associated proteins. Levels of SMC-derived proteins such as αSMA, however, were reduced by the filtration step, consistent with imaging data showing a reduction in larger diameter αSMA+ vessels within the output material from this approach. These observations support the notion that the enrichment and filtration techniques presented herein yield samples more likely to represent a larger proportion of BBB-associated capillaries with modest to lower inclusion of non-capillary sized vessels.

**Figure 3.**
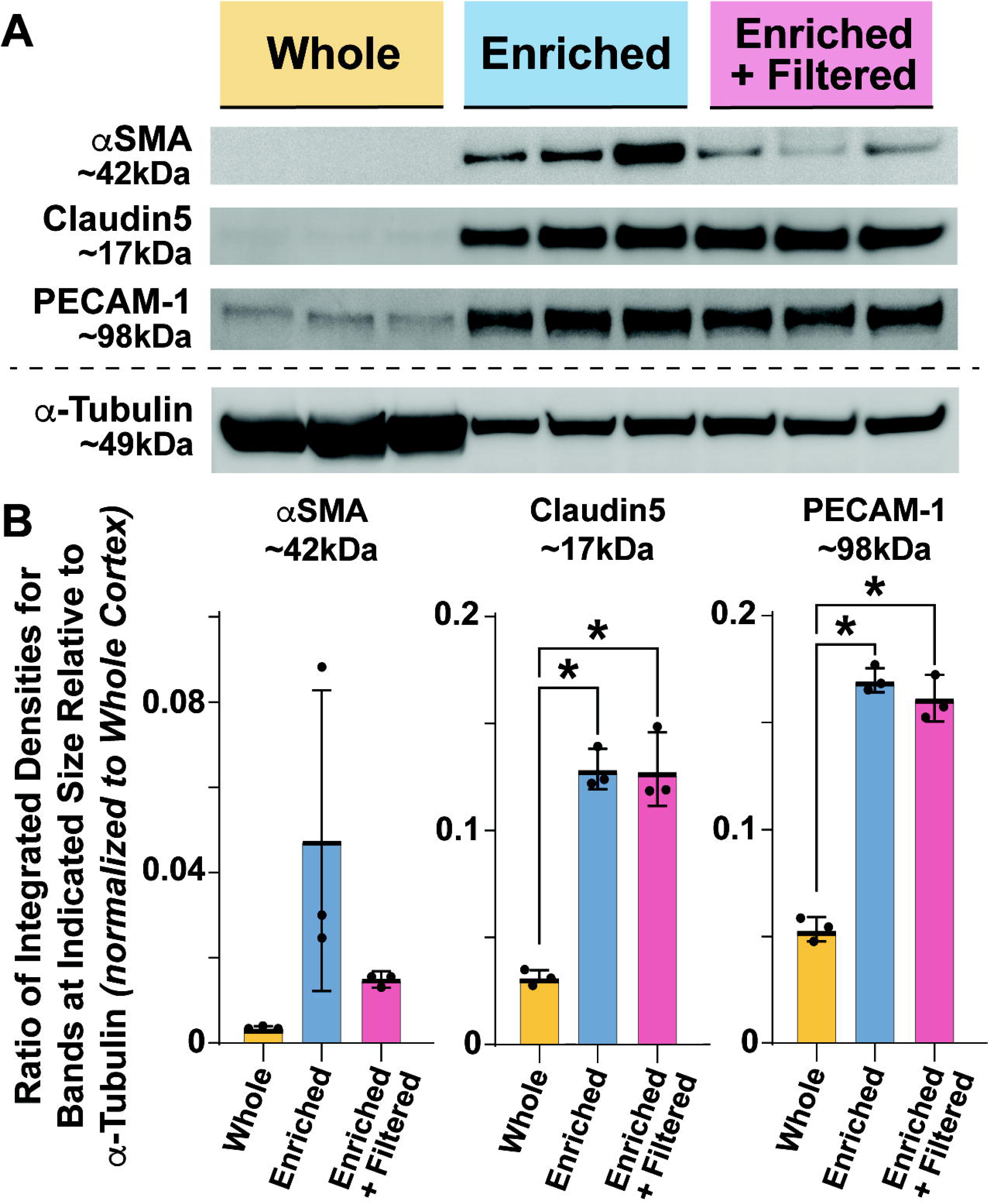
Vascular-Related Proteins are Enriched Following Density Gradient Centrifugation of Mechanically Disaggregated Murine Brain with Additional Filtration Limiting Contributions from Vascular SMCs. (A) Representative western blot images of αSMA (∼42kDa), Claudin5 (∼17kDa), and PECAM-1 (∼98kDa) with α- tubulin (∼49kDa) shown for reference. Protein lysates were collected from the output of each protocol: Group 1—Whole Cortex (lanes 1-3), Group 2—Enriched by DGC (lanes 4-6), and Group 3—Enriched by DGC and Filtered (lanes 7-9). (B) Graph of the ratios of integrated densities for bands at the indicated sizes relative to α-tubulin. Measurements were taken from the Whole Cortex group (Group 1), shown as individual data points with the average shown by the yellow bar. Measurements taken from Groups 2 (Enriched by DGC) and 3 (Enriched by DGC and filtered) were normalized to the Whole Cortex average for relative comparison across the different protocol groups. Individual data points are shown with averages represented by bars, blue for Group 2 (Enriched) and pink for Group 3 (Enriched+Filtered). Error bars are standard deviation, and * denotes p ≤0.05 by ordinary one-way ANOVA followed by unpaired Tukey’s test comparisons for the groups indicated. Four (4) brain samples were pooled for each biological replicate, with n=3 biological replicates per protocol output. Original blots are provided in Supplemental Figures 1 and 2.

### Transcriptional Profiling of Post-Collection Samples Confirms Filtration as a Means to Lower Inclusion of Non-Capillary Vessels in Output Material

As an orthogonal approach to analyze the content from each vessel isolation protocol, we selected gene expression targets based on vascular network hierarchy and applied them to output samples from which mRNA was collected. Complementing our morphology- and protein-based analyses, we measured *Acta2* (αSMA) transcripts from whole murine cortex, samples enriched by DGC, and those that were enriched and filtered (Figure 4A). Enrichment increased the levels of *Acta2*/αSMA mRNA around 4- fold in output samples as compared to whole cortex, consistent with a high level of non-capillary vessels associated with SMCs. Filtering enriched samples reduced these transcripts substantially but did not completely remove them (Figure 4A), as seen in residual SMC contributions to the imaging and protein data (Figures 2 & 3). This same trend was observed when measuring other SMC-related genes, particularly those associated with arteries and arterioles, such as myosin heavy chain protein 11 (*Myh11*, Figure 4B) and calponin 1 (*Cnn1*, Supplementary Figure 3).

**Figure 4.**
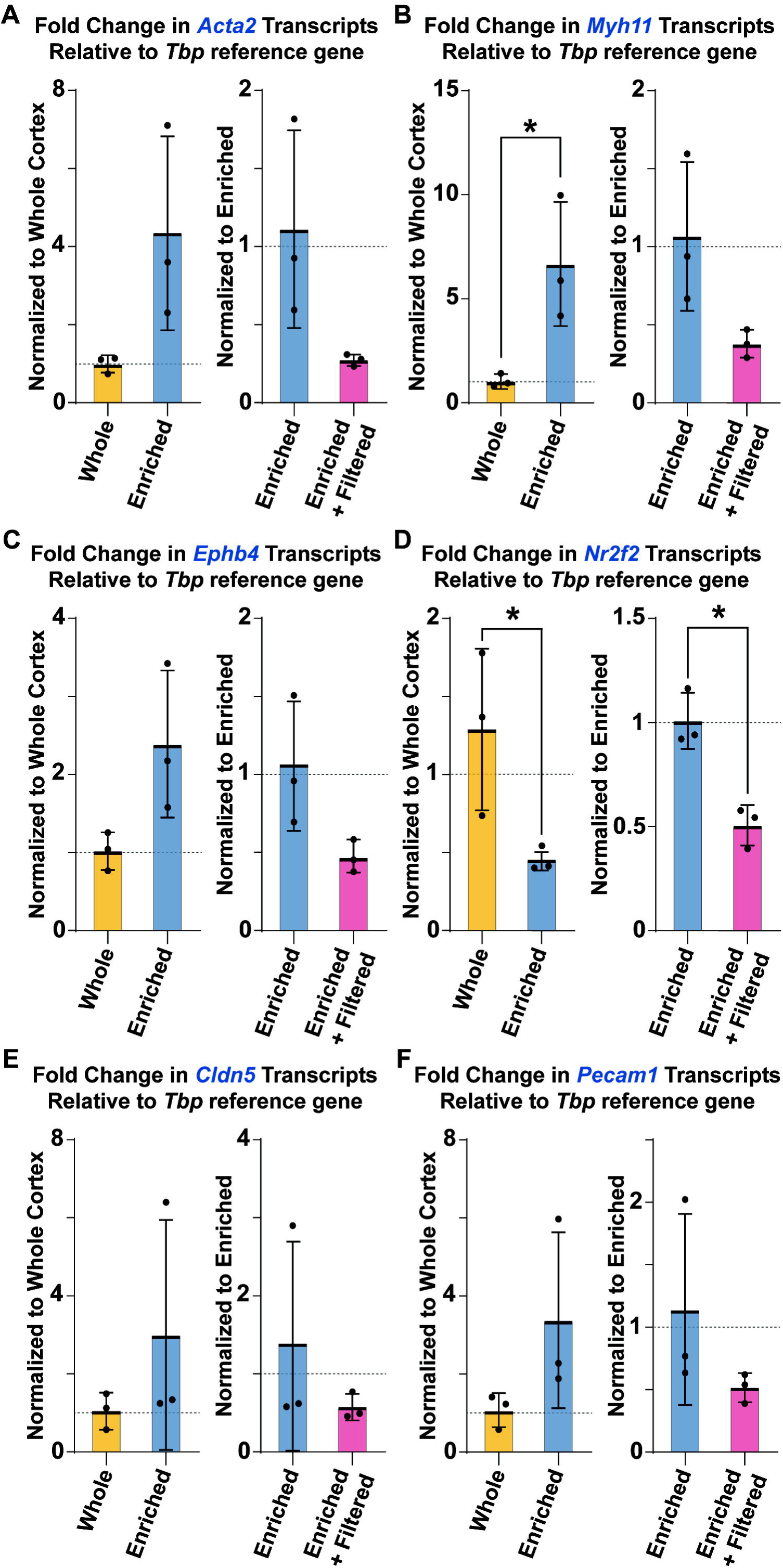
Gene Expression Analysis of Output Material from each Protocol Underscores the Utility of Filtration in Reducing Non-Capillary Vessels. For all mRNA transcripts evaluated using qRT-PCR, *Tbp* was our housekeeping gene, and we normalized the measurements for each protocol group separately to better appreciate the fold difference in transcript levels between them. Specifically, enriched samples (Group 2) were referenced to whole cortex samples (Group 1) (left graph), and enriched and filtered samples (Group 3) were referenced to enriched samples (Group 2) (right graph). Dotted lines indicate “1” for the reference sample. Individual data points are shown with averages represented by bars – Whole Cortex (Group 1): yellow bars, Enriched by DGC (Group 2): blue bars, and Enriched and Filtered (Group 3): pink bars. Relative quantitation was calculated for the following genes: (A) *Acta2,* (B) *Myh11,* (C) *Ephb4,* (D) *Nr2f2,* (E) *Cldn5,* and (F) *Pecam1*. Error bars are standard deviation, and * denotes p≤0.05 by Student’s T-test. n=3 biological replicates per protocol output. Additional supporting qRT-PCR data provided in Supplemental Figure 3.

Based on these observations, we further hypothesized that inclusion of larger diameter vessels within venous regions of the cerebrovasculature would mirror that of arterial segments. To test this idea, we selected several EC markers that have been strongly associated with a venous identity. Specifically, EphB4 has emerged as a key factor in the specification of venous ECs^31^ Quantifying *Ephb4* transcripts revealed a 2.5-fold increase within DGC samples without filtration (Figure 4C), suggesting venous regions of brain vascular networks could be enriched with this approach. Filtering samples following enrichment reduced *Ephb4* mRNA by about half, reflecting a similar decrease in SMC markers tested and suggesting a partial removal of veins/venules with follow-on filtration (Figure 4C). In addition, we measured the levels of a transcript encoding another well-accepted marker of venous ECs, particularly in the developmental setting — chicken ovalbumin upstream promoter transcription factor-II (COUP-TFII, or Nuclear Receptor 2F2, gene name: *Nr2f2*)^32,33^. Interestingly, these mRNA transcripts were significantly reduced with DGC, and their levels were lowered further with subsequent filtration (Figure 4D). From a purely vascular biology perspective, these results were initially unexpected. However, in considering recent insights from neuroscience studies regarding COUP-TFII/*Nr2f2* expression in neuronal subtypes^34,35^, our observations align with a substantial depletion of these neurons by DGC with an additional reduction of these cells by subsequent filtration, potentially alongside the removal of venous ECs.

To further compare the effect of different isolation strategies on vascular-associated material collected, we selected the more general EC markers utilized in our imaging and western blot analyses, specifically Claudin-5/*Cldn5* and PECAM-1/*Pecam1.* Enrichment of vessels by DGC increased the relative transcript levels for each of these targets compared to whole cortex preparations (Figure 4E-F), reflecting a similar trend seen in protein levels for these markers (Figure 2). In contrast with our western blot observations, we found that DGC plus filtration yielded lower levels of these markers relative to DGC alone (Figure 4E-F). While the protein and mRNA levels for a given target may not align 1:1 for various biological reasons, this discrepancy may also be capturing important differences between ECs in larger diameter vessels and those within microvessels. Morphological differences are known to exist between these two populations, which may in turn relate to how EC junctions are regulated in cerebral arteries and veins compared to capillaries.

## DISCUSSION

Capillaries that compose the blood-brain barrier (BBB) exist in a continuum within cerebrovascular networks, bridging upstream arterioles/arteries with downstream venules/veins and facilitating selective exchange. Because this unique vascular region plays such a vital role in neurological health and disease, it has received significant attention in the scientific community, inspiring the development of various methods to separate BBB capillaries from (i) adjacent non-vascular cells and (ii) flanking non-capillary vessels. Comprehensive characterization of the output material from these protocols is critical in providing the most accurate context for follow-on experiments, data analysis, and associated interpretations and conclusions. Here, visual assessment of output samples using several fluorescent labels revealed a substantial reduction in non-vascular tissue following DGC and filtration. This approach also enriched capillary-sized vessels but could not fully exclude microvascular segments associated with αSMA+ vSMCs. Analysis of protein content by western blot further highlighted the enrichment of vascular markers in samples processed by DGC, with additional filtration maintaining BBB-associated markers (e.g. Cldn5) and reducing – though not fully removing – arterial/venous markers such as αSMA. Transcriptional profiling of arterial and venous vSMC and EC markers largely reflected these trends of DGC plus filtration yielding samples with reduced non-vascular and non-capillary contributions. Applying the lens of vascular network hierarchy to the post-collection analysis provides a more complete assessment of the material yielded from this optimized BBB enrichment protocol. This approach is critical for providing a more accurate representation of the specific cerebrovascular structures being used in downstream applications, the interpretation of follow-on observations, and the assignment of BBB properties to specific regions of the brain vasculature.

Current and emerging methodologies designed to isolate and enrich cerebrovascular fractions, and specifically BBB microvessels, will invariably contain certain levels of non-vascular tissue/cells as well as vessels not within capillary networks. It is essential therefore to rigorously evaluate the collection output and interpret all associated data within the context of this assessment. Numerous studies include analysis of BBB-associated transcriptional markers and morphological characteristics^23–25^, but some EC junction markers suggested to strictly define the BBB are also found in ECs of cerebral arteries/arterioles, veins/venules^28–30^, and non-BBB vasculature (i.e. circumventricular organs, choroid plexus, dura mater)^36^ Though challenges exist with sufficient sampling and accuracy of high throughput methods, establishing the physical attributes of collected vessels can be a useful complementary approach. We found that tissue aggregation may introduce another complication with imaging-only strategies, as clear discernment of larger vessels from microvessels hampered efforts to quantify each population. Therefore, extending this analysis to detect the presence of vSMC markers such as αSMA, as done herein and in other studies^21,26^, provides another layer of insight into the population of vessels yielded by a particular method. We found that protein and transcriptional profiling of arterial and venous markers, which is often limited or absent from BBB-focused studies, represents another important step towards comprehensive evaluation of large-vessel content yielded by cerebrovascular enrichment protocols. This strategy may be a useful addition to studies aimed at subsequent culture of brain vascular cells where arterial and/or venous vSMCs may be present alongside BBB- associated pericytes^37^ Because these two cell types share expression of several overlapping markers (e.g. *Cspg4*/NG2, *Pdgfrb*), this assessment scheme may therefore aid in efforts to distinguish pericyte and vSMC behaviors in certain culture environments and with specific interventions.

Density gradient centrifugation (DGC) and follow-on filtration techniques have emerged as promising steps in protocols designed to yield capillary-sized vessels from mechanically and/or enzymatically dissociated brain tissue^21–27^. Here, we drew from these previous approaches to optimize an enrichment strategy that was complemented by a focus on vascular network hierarchy in the assessment of output material. We preferred mechanical disaggregation of the cerebrovasculature, as several studies have suggested enzymatic methods may compromise mRNA quality^38^ and inadvertently activate certain cell types^39^ In addition, the optimized protocol presented herein did not require: (i) the use of antibody labeling, which may be cost prohibitive and may further promote cellular activation^40^ or (ii) specific transgenic animals that may complicate data interpretation^41^ Previous studies utilizing these approaches have also indicated that protein post-translational modifications (e.g. phosphorylation, ubiquitination, glycosylation) are retained, allowing evaluation of signaling pathway activity, among other downstream analyses^27^. We also found that pooling brain samples was necessary to yield sufficient levels of output material for certain applications; however, our input numbers were likely lower than the amount required for higher-purity/lower-yield approaches such as ultra-speed centrifugation with additional filtration steps and cell sorting (MACS, FACS).

As with all scientific studies and protocols, key limitations must be acknowledged. Here, we need to consider all data alongside the fact that these enriched cerebrovascular fragments have been removed from their native environment and are likely affected to some degree by this process. Studies employing these techniques should therefore incorporate orthogonal experiments and analyses to validate data from BBB-targeted isolates. In addition, we focused our post-collection assessment on a subset of vSMC markers and arterial/venous EC signatures. Future studies extending this work could include a broader array of genes predominantly expressed by vSMCs, perhaps distinguishing those with pericyte-associated transcripts where possible. It might also be of interest to more extensively profile non-vascular cells (i.e. astrocytes, microglia, neurons) with the methodology applied here, to gain a better appreciation for the relative ratios of each cellular compartment in the output material. Lastly, we were intrigued by observing Claudin5 and PECAM-1 protein remaining at relatively similar levels between the enrichment and enrichment plus filtration groups, yet the number of transcripts for these markers appeared to decrease with filtration, although not to a statistically significant degree. The current study was not designed to address this specific observation. Thus, we can only speculate that this mismatch may be due to inherent morphological and/or transcriptional distinctions between capillary ECs and those within larger diameter vessels. These differences may in fact reflect the nature of EC junction regulation in cerebral capillaries relative to arteries/veins, but this idea would need to be confirmed with follow-up studies with this as the primary focus.

The protocols and analyses presented herein can be applied to many future applications focused on rigorously assessing BBB changes during specific pathologies, interventions, or experimental conditions. For instance, Alzheimer’s Disease onset and progression likely involve elements of maladaptive BBB remodeling that are still being fully resolved^42^. At the same time, effective delivery of therapeutic agents to treat this devastating condition will also depend on understanding drug transit into the brain parenchyma at these sites of blood-tissue exchange i.e. the cerebral capillaries.

Fluctuations in metabolic inputs (e.g. glucose in diabetes) or in hemodynamic forces (e.g. reperfusion following stroke intervention) are additional factors that may alter BBB integrity and impact outcomes in the clinic and in pre-clinical models^11^. These methods can therefore be incorporated across a broad range of important BBB-focused studies.

## Supplemental Information

Supplemental information can be found online and includes original western blot images and additional qRT-PCR analysis.

## Acknowledgements

We would like to thank all of the members of the Chappell lab for their support and valued assistance on this project, both materially and intellectually.

This work was supported in part by funding from the National Institutes of Health (R01HL159512 to J.W.S., R01HL146596 to J.C.C., R01NS105807 to S.R., F31HL168946 to H.A.) and the American Heart Association (AHA, 19TPA34910121 to J.C.C. and 23PRE1025483 to K.L.Y.).

## Author Contributions

H.A., C.M.B., K.L.Y., and J.C.C. conceptualized and optimized methodology. H.A. performed experiments. H.A. and R.H. analyzed datasets. H.A. and J.C.C. wrote the manuscript. H.A., A.B., and J.S. provided critical data for the manuscript. S.R., J.W.S., S.L., and J.C.C. acquired financial support. All authors have reviewed and approved the final version of this manuscript.

## Inclusion and Diversity

We support inclusive, diverse, and equitable conduct of research.

## Abbreviations

*EC – Endothelial Cell, ECM – Extracellular Matrix, vSMC – vascular Smooth Muscle Cell, PC – Pericyte, BBB – Blood-Brain Barrier, MACS – Magnetic- Activated Cell Sorting, FACS – Fluorescence-Activated Cell Sorting, DGC – Density Gradient Centrifugation, BSA – Bovine Serum Albumin, α-SMA – α-smooth muscle actin, PECAM-1 – Platelet-Endothelial Cell Adhesion Molecule-1, PDGFRβ – Platelet- Derived Growth Factor Receptor-β*

## Research Topics

CP: Neuroscience

## STARMETHODS

### KEY RESOURCES

| REAGENT or RESOURCE | SOURCE | IDENTIFIER / CATALOG # |
| --- | --- | --- |
| **Antibodies** |  |  |
| Polyclonal Goat IgG Anti-mouse PECAM-1/CD31 | Biotechne | Cat. #AF3628 |
| Monoclonal Mouse IgG Anti-mouse αSMA | Millipore / Sigma | Cat. #A5228 |
| Monoclonal Mouse IgG Anti-mouse αSMA, FITC Conjugate | Millipore / Sigma | Cat. #F3777 |
| Polyclonal Rabbit IgG Anti-mouse Claudin5 | Invitrogen | Cat. #34-1600 |
| Polyclonal Goat IgG Anti-mouse PDGF Receptor-β | Biotechne | Cat. #AF1042 |
| Polyclonal Rat IgG Anti-mouse PDGF Receptor-β | Invitrogen | Cat. #14-1402-82 |
| Monoclonal Mouse IgG Anti-mouse α-tubulin | Millipore / Sigma | Cat. #T6199 |
| Polyclonal Donkey IgG Anti-mouse AlexaFluor568 | Invitrogen | Cat. #A10037 |
| Polyclonal Donkey IgG Anti-goat AlexaFluor647 | Jackson ImmunoResearch | Cat. #705-605-147 |
| Polyclonal Donkey IgG Anti-rabbit AlexaFluor647 | Jackson ImmunoResearch | Cat. #711-605-152 |
| Polyclonal Donkey IgG Anti-rat DyLight550 | Invitrogen | Cat. #SA5-10027 |
| Isolectin GS-IB4 | Invitrogen | Cat. #132450 |
| **Chemicals, peptides, and recombinant proteins** |  |  |
| HEPES, 1M | ThermoFisher Scientific | Cat. #15630106 |
| HBSS | ThermoFisher Scientific | Cat. #14170112 |
| Dextran | Millipore / Sigma | Cat. #31390-25G |
| BSA | Millipore / Sigma | Cat. #126609-10GM |
| RIPA Solution | ThermoFisher Scientific | Cat. #89901 |
| Halt™ Protease and Phosphatase Inhibitor Cocktail (100X) | ThermoFisher Scientific | Cat. #78442 |
| Chloroform | Fisher Scientific | Cat. #ICN19400225 |
| Trizol | Invitrogen | Cat. #15596018 |
| Ethanol, Absolute (200 Proof), Molecular Biology Grade | Fisher Scientific | Cat. #BP2818500 |
| 2-Propanol | Fisher Scientific | Cat. #416-4 |
| EDTA (100x) | ThermoFisher Scientific | Cat. #1860851 |
| **Experimental models: Organisms/strains** |  |  |
| Mouse: C57BL/6J | The Jackson Laboratory | RRID: IMSR_JAX:000664 |
| Mouse: *Cspg4:DsRed/+* (*NG2:DsRed/+*) | The Jackson Laboratory | RRID: IMSR_JAX:008241 |
| **Software** |  |  |
| ImageJ2 / FIJI |  |  |
| ImageLab v5.2.1 software | Bio-Rad |  |
| **Other** |  |  |
| KIMBLE Dounce tissue grinder set | Millipore / Sigma | Cat. #D9063 |
| EZFlow® Cell Strainer, 100µm | Foxx Life Sciences | Cat. #410-0003-OEM |
| Filter paper (Whatman) | Millipore / Sigma | Cat. #WHA1114125 |
| Low Binding Tips | Genessee Scientific | Cat. #23165RS |

### RESOURCE AVAILABILITY

#### Lead contact

Further information and requests for resources and reagents should be directed to and will be fulfilled by the lead contact, Dr. John C. Chappell.

#### Materials availability

This study did not generate any new unique reagents.

### EXPERIMENTAL MODEL AND SUBJECT DETAILS

#### In vivo animal studies

In the current study, we utilized brain tissue samples from C57BL/6 and transgenic mice harboring a gene construct labeling microvascular pericytes as well as other cell types expressing *Ng2/Cspg4* (e.g. oligodendrocyte precursors), specifically with the DsRed2 fluorophore (STOCK Tg(Cspg4-DsRed.T1)1Akik/J, NG2DsRedBAC, The Jackson Laboratory, Strain #008241). In total, 70 adult mice were used for this study – a mix of males and females between the ages of 3- and 9-months old, under approval granted by the Institutional Animal Care and Use Committee (IACUC), via protocol #23-167. All experiments involving animal use were performed following review and approval from the IACUC at Virginia Tech. All experimental protocols were reviewed and approved by Virginia Tech Veterinary Staff and the IACUC. The Virginia Tech NIH/PHS Animal Welfare Assurance Number is A-3208-01 (Expires: 31 July 2025).

### METHODS DETAILS

#### Microvascular Isolation

See Figure 1 for a broad overview schematic of the experimental workflow. To enhance reproducibility, the following is the stepwise protocol used in this study:

1. Prepare isolation vessel solutions and place on ice: **Solution 1**: add 1.5 ml of HEPES 1M to 150 ml of HBSS **Solution 2**: add 3.6 g of Dextran to 20 ml of **Solution 1 Solution 3**: add 1 g of BSA to 100 ml of **Solution 1** (1%)
2. Gather the following essential materials:

a. 15 ml conical tubes
b. 50 ml conical tubes
c. 2 ml centrifuge tube
d. Buckets for ice
e. 7-ml Dounce tissue grinder
f. 100 µm-mesh filter (cell strainer)
g. Petri dish
h. Blotting (filter) paper
i. Automated Shaker
j. Kimwipes®
3. Humanely euthanize animal according to approved methods and then perfuse cold PBS via an intracardiac catheter *(Note: perfusion may be optional depending on downstream application.)*.
4. Carefully extract the brain and roll it gently on blotting paper to remove the meninges and any meningeal vessels.
5. Immediately transfer it to a Petri dish containing ice-cold **Solution 1**; rinse the brain in this solution and ensure that the entire brain remains submerged in this cold solution.
6. Carefully remove the cerebellum and olfactory bulb, and then isolate the cortices.
7. Homogenize the cortical tissue using a loose-fit, 7-ml Dounce tissue grinder (see Key Resources table). *Important*—*Keep grinder submerged in ice during homogenization.* Perform an initial 10 strokes in 1 ml of ice-cold **Solution 1** at a steady pace, *<u>without</u>* twisting the pestle. Add 7 ml of **Solution 1** to the homogenate, and perform an additional five strokes in a total of 8 ml.
8. Pour the homogenized tissue into a 15 ml conical tube. *Note: If processing multiple samples, take care to adjust the volumes of the tubes with additional **Solution 1** as needed, so that all samples contain the same amount of mixture*.
9. Centrifuge at 2,000×g for 10 min at 4°C.
10. Pour out the supernatant and carefully rest the inverted tube on Kimwipe to absorb excess medium/debris, but be sure to not disturb the pellet.
11. Resuspend the pellet in 10 ml of **Solution 2**, and then shake vigorously for 1 min.
12. Centrifuge at 4,400×g for 15 min at 4°C. *Note: If the protocol is trending towards success, a reddish pellet will appear; however, the pellet color will be clearer if perfusion was done*.
13. Remove the top layer and the supernatant carefully to avoid contaminating the pellet with debris, and then clean the inside surfaces of the conical tube with a Kimwipe®.

*Note: If enrichment of all sizes of cerebral vessels are needed, then proceed to Step 18. However, if microvessel enrichment is desired, continue to the following filtration steps*.

14. Resuspend the pellet with 1 ml of **Solution 3** using low-binding tips. *It is important to wet the tips slightly before use in **Solution 3**.* Be sure to disrupt the pellet completely.
15. Add 5 ml of **Solution 3**, pipetting up and down several times.

The vessels and solution from the previous step can be filtered on a 100 µm-pore mesh cell strainer. We found that larger vessels are retained on top of the filter, while the flow- through contains predominantly microvasculature.

16. Place a 100 µm-mesh filter on a 50 ml conical tube, and then activate the strainer with ice cold **Solution 3** to equilibrate and allow for more efficient filtration.
17. Pour the vessel preparation from the previous step onto the filter and rinse the tube and membrane with 15 ml of ice-cold **Solution 3**.
18. Centrifuge the conical tube at 4400×g for 5 min at 4°C.
19. Remove the supernatant, and then resuspend the pellet in 1 ml of ice-cold **Solution 3.**
20. To further concentrate the output material, add the resuspended pellet to a 2 ml microcentrifuge tube, and then centrifuge for 5 min at 2000×g at 4°C.
21. Discard the supernatant.

Note: For Downstream applications, we recommend pooling brain samples to increase the overall yield. For a reasonable yield of mRNA, we recommend pooling a minimum of 2 brains. For protein collection, we recommend a minimum of 4 brains.

#### Immunostaining and Confocal Imaging

For immunostaining isolated vessels, we recommend using Low Binding tips in all the steps. Resuspend the vessels gently after every incubation or wash. To remove staining or wash solutions, centrifuge for 1 min at 1000×g at 4°C. The base staining solution consisted of PBS with 0.5% Triton-X 100 (PBS-T), and all primaries and secondaries were diluted in this PBS-T solution.

1. Following the last step of the microvascular isolation protocol, suspend the pellet in 250 µl of 4% paraformaldehyde (PFA) and allow to incubate for 30 min to 1 hr at room temperature (∼25°C). *Note: PFA is a carcinogen and should be handled with extreme caution*.
2. Carefully, centrifuge (1 min at 1000×g at 4°C) and then discard the 4% PFA in the appropriate waste stream.
3. Resuspend the pellet with 500 µl of 1x PBS, and incubate for 5 min at room temperature, and then centrifuge (1 min at 1000×g at 4°C). ***Repeat this step 3 times***.
4. After the last wash, add blocking solution (PBS-T + 5% BSA) and then resuspend the vessels gently.
5. Incubate in PBS-T for 1 hr at room temperature, centrifuge (1 min at 1000×g at 4°C), and discard the supernatant blocking solution.
6. Add primary antibodies at appropriate concentrations (PECAM-1/CD31: 1:200, PDGFRβ: 1:200, αSMA-FITC: 1:200), and incubate for 3 hours at room temperature, or leave overnight at 4°C.
7. Wash with PBS at room temperature. **Repeat 3 times.**
8. Add secondary antibodies (all at 1:500) in PBS-T for 3 hrs at room temperature, or leave overnight at 4°C.
9. If desired, stain cell nuclei with DAPI (1:1000 in PBS-T) for 30 min at room temperature.
10. After removing the DAPI solution, resuspend the vessels in 100 µl of PBS.
11. For mounting, outline a glass microscope slide using a wax or hydrophobic pen and then let it dry.
12. Gently add your resuspended sample to the slide, ensuring that the sample is distributed within the outline and allow fragments to settle onto the surface of the slide.
13. Carefully remove any excess PBS using Kimwipes®, and let it dry for 10-20 minutes.
14. Apply 1-2 drops of mounting media, and then cover with a glass coverslip, starting at an angle and allowing it lower gently onto the sample.
15. Let the glass coverslip settle, and the mounting medium distribute for 1 hr in the dark at room temperature.
16. Seal the sides of the glass coverslip with clear nail polish and let dry.
17. Store slide at 4°C until ready for imaging.

Here, we imaged on a Zeiss LSM 880 point-scanning laser confocal microscope using 10×, 20×, and 40× objectives. Parameters were adjusted to avoid signal saturation, and images were taken through the thickness of samples along the z-axis. These “z-stack” images were then compressed into a single image.

#### Western Blot

Following the last step of the microvascular isolation protocol, add the appropriate amount of RIPA to the sample and then “snap freeze” by placing on dry ice for several minutes. Store immediately at -80°C overnight or until used. Depending on tissue size, 100 to 300 uL of RIPA buffer will be needed per sample. In general, a “working RIPA solution” included 970 μl of commercially available 1x RIPA buffer with 30 μl of 100x protease inhibitor cocktail (HALT^TM^) and 2 μl of EDTA solution. The general western blotting procedure was as follows:

1. Remove samples from -80°C and immediately place on ice to thaw. *Note: Always keep your protein lysate on ice to minimize protein degradation*.
2. Homogenize the tissue using 20-, 23-, and 25-gauge needles consecutively until the sample is fully homogenized in working RIPA solution.
3. Once homogenization is complete, incubate tubes on ice for 20 minutes. Vortex tubes several times during the incubation to ensure even mixing.
4. Centrifuge tubes for 20 min at 15,000×g at 4°C.
5. Collect the supernatant and transfer to a new 1.5 mL Eppendorf tube kept on ice.
6. Depending on the volume of protein lysate and the desired concentration, small aliquots at the appropriate volume/concentration can be generated and frozen to avoid repeated freeze/thaw of stock protein solution. If aliquots are made, snap freeze on dry ice and store at -80°C.
7. For samples being processed for western blot, perform protein quantification using a standard assay such as the Bradford Protein Assay (BioRad), and normalize samples for SDS-PAGE in LDS sample buffer (ThermoFisher Scientific).
8. Load and separate proteins using standard gel electrophoresis techniques. Here, we separated proteins using Invitrogen Bolt Bis-Tris Plus 4-12% Mini Protein gels with corresponding Invitrogen reagents and buffers. Be sure to include a protein ladder for size analysis. Here, we included the Invitrogen SeeBlue Plus 2 Protein Standard and allowed for a 26-minute protein migration at 200V.
9. Transfer protein to an appropriate membrane for immunostaining. Here, protein staining was performed on BioRad PVDF mini membrane after transfer, using the “Mixed Molecular Weight (MW)” program on a BioRad TransBlot Turbo®.
10. Primary antibodies should be diluted in TBS and incubated overnight at 4°C (PECAM-1/CD31: 0.4 ug/mL; Claudin5: 0.25 ug/mL; αSMA: 2.3 ug/mL; α-tubulin: 0.2 ug/mL), while secondary antibodies diluted in TBS can be incubated for 2 hours at room temperature (donkey anti-goat PECAM-1/CD31: 3 ug/mL; donkey anti-rabbit Claudin5: 1.5 ug/mL; donkey anti-mouse αSMA: 2 ug/mL; donkey anti- mouse α-tubulin: 0.4 ug/mL).
11. Image on an appropriate blot imaging system such as a BioRad ChemiDoc®.
12. After imaging, membranes can be stripped using 1x Millipore ReBlot Plus® Strong Antibody Stripping Solution (Millipore/Sigma, Cat. #2504 for 10x), washed with double distilled water and TBS, and re-stained as needed.

#### Quantitative Real-Time Polymerase Chain Reaction (qRT-PCR)

Following the last step of the microvessel isolation protocol, add 500 ml of Trizol to the sample and store immediately in -80°C overnight. To begin RNA extraction, gather the following materials: Trizol or Trizol LS reagent, Chloroform, Isopropanol (2-propanol), 75% Ethanol in RNase-free water (be sure to make this fresh). *Note: all centrifugation steps are done at 4°C*. To isolate RNA:

1. Homogenize the sample in Trizol using 20-, 23-, and 25-gauge needles sequentially until the sample is fully lysed.
2. Incubate samples for 5 minutes in Trizol at room temperature.
3. Add 100 ul of chloroform per 500 ul of Trizol and shake vigorously for 20 sec. Incubate 3 mins at room temperature. *Note: Perform these steps with extreme caution, as Trizol and chloroform are caustic and very hazardous*.
4. Centrifuge at 12,000×g for 15 min at 4°C.
5. Transfer top, clear aqueous phase containing RNA into new tube making sure *<u>not</u>* to disturb the white DNA layer beneath the RNA layer.
6. Add 250 μl of isopropanol per 500 μl Trizol to the aqueous phase and gently mix.
7. Incubate tubes for 10 min at room temperature.
8. Spin at 12,000×g for 10 min at 4°C. Note: with low yield samples, the RNA pellet may be very difficult to visually identify. Position the tube in the centrifuge with the back hinge of the cap pointing out to indicate where the pellet will collect (i.e. on the backside of the tube).
9. Carefully remove the isopropanol *<u>without</u>* disturbing the RNA pellet. This may be achieved by either pipetting it out or pouring it off.
10. Re-suspend the pellet with 500 μl of 75% ethanol per 500 μl Trizol. *Note: Ensure that the pellet does not get stuck inside the pipette tip*.
11. Centrifuge at 12,000×g for 10 min at 4°C.
12. *<u>Very gently</u>*, invert the tube over onto a dry Kimwipe® that was previously soaked with RNase Displace^TM^.
13. Leave the tube inverted to air dry on the Kimwipe® for 10 min at room temperature.
14. After it dries completely, re-suspend immediately in RNase-free water (∼10-20μl).
15. Once re-suspended, place the samples on ice.
16. Use a Nanodrop® to measure concentration and purity. Store RNA at -80C
17. qRT-PCR can be performed by many standard protocols. Here, each sample was run in triplicate on a QuantStudio Flex 6. Applied Biosystem Taqman assay probes and Master Mix were used on 96- and 384-well optical plates.

### QUANTIFICATION AND STATISTICAL ANALYSIS

GraphPad Prism 8 software was used for statistical analysis. For measurements where statistical comparisons are shown, we applied a Student’s t-test or an ordinary one-way ANOVA followed by unpaired Tukey’s test comparing individual groups. Statistical significance was set at a P value less than or equal to 0.05. Measurements were taken across a minimum of n=3-4 biological replicates for each protocol group, with technical replicate measurements taken where possible. Biological replicates were individual brain samples (whole cortex groups) or pooled brain samples for the enrichment protocols (2 for each replicate to generate sufficient mRNA, and 4 for each replicate to yield material for protein analysis).

### ADDITIONAL RESOURCES

None.

## SUPPLEMENTAL INFORMATION

**Supplemental Figures 1 and 2.** Original western blots in support of Figure 3.

**Supplemental Figure 3.** Calponin1 (*Cnn1*) Expression Analysis of Output Material from each Protocol Underscores the Utility of Filtration in Reducing Non-Capillary Vessels. *Cnn1* transcripts were evaluated using qRT-PCR. *Tbp* was our housekeeping gene, and we normalized the measurements for each protocol group separately to better appreciate the fold difference in transcript levels between them. Specifically, enriched samples (Group 2) were referenced to whole cortex samples (Group 1) (left graph), and enriched and filtered samples (Group 3) were referenced to enriched samples (Group 2) (right graph). Dotted lines indicate “1” for the reference sample. Individual data points are shown with averages represented by bars – Whole Cortex (Group 1): yellow bars, Enriched by DGC (Group 2): blue bars, and Enriched and Filtered (Group 3): pink bars. Error bars are standard deviation, and * denotes p≤0.05 by Student’s T-test.

## REFERENCES

1. Trimm, E., and Red-Horse, K. (2023). Vascular endothelial cell development and diversity. Nat Rev Cardiol 20, 197–210. 10.1038/s41569-022-00770-1.

2. Wagenseil, J.E., and Mecham, R.P. (2009). Vascular Extracellular Matrix and Arterial Mechanics. Physiol Rev 89, 957–989. 10.1152/physrev.00041.2008.

3. Hennigs, J.K., Matuszcak, C., Trepel, M., and Körbelin, J. (2021). Vascular Endothelial Cells: Heterogeneity and Targeting Approaches. Cells 10, 2712. 10.3390/cells10102712.

4. Potente, M., and Makinen, T. (2017). Vascular heterogeneity and specialization in development and disease. Nat Rev Mol Cell Biol 18, 477–494. 10.1038/nrm.2017.36.

5. Fish, J.E., and Wythe, J.D. (2015). The molecular regulation of arteriovenous specification and maintenance. Developmental Dynamics 244, 391–409. 10.1002/dvdy.24252.

6. Aird, W.C. (2007). Phenotypic heterogeneity of the endothelium: I. Structure, function, and mechanisms. Circ Res 100, 158–173. 100/2/158 [pii] 10.1161/01.RES.0000255691.76142.4a.

7. Aird, W.C. (2007). Phenotypic heterogeneity of the endothelium: II. Representative vascular beds. Circ Res 100, 174–190. 100/2/174 [pii] 10.1161/01.RES.0000255690.03436.ae.

8. Chavkin, N.W., and Hirschi, K.K. (2020). Single Cell Analysis in Vascular Biology. Front Cardiovasc Med 7. 10.3389/fcvm.2020.00042.

9. Gifre-Renom, L., Daems, M., Luttun, A., and Jones, E.A. V. (2022). Organ-Specific Endothelial Cell Differentiation and Impact of Microenvironmental Cues on Endothelial Heterogeneity. Int J Mol Sci 23, 1477. 10.3390/ijms23031477.

10. Ribatti, D., Nico, B., Crivellato, E., and Artico, M. (2006). Development of the blood-brain barrier: A historical point of view. The Anatomical Record Part B: The New Anatomist 289B, 3–8. 10.1002/ar.b.20087.

11. Hoque, M.M., Abdelazim, H., Jenkins-Houk, C., Wright, D., Patel, B.M., and Chappell, J.C. (2021). The cerebral microvasculature: Basic and clinical perspectives on stroke and glioma. Microcirculation 28. 10.1111/micc.12671.

12. Mastorakos, P., and McGavern, D. (2019). The anatomy and immunology of vasculature in the central nervous system. Sci Immunol 4. 10.1126/sciimmunol.aav0492.

13. Bell, B., Anzi, S., Sasson, E., and Ben-Zvi, A. (2023). Unique features of the arterial blood–brain barrier. Fluids Barriers CNS 20, 51. 10.1186/s12987-023-00450-3.

14. Siletti, K., Hodge, R., Mossi Albiach, A., Lee, K.W., Ding, S.-L., Hu, L., Lönnerberg, P., Bakken, T., Casper, T., Clark, M., et al. (2023). Transcriptomic diversity of cell types across the adult human brain. Science (1979) 382. 10.1126/science.add7046.

15. Saliba, A.-E., Westermann, A.J., Gorski, S.A., and Vogel, J. (2014). Single-cell RNA-seq: advances and future challenges. Nucleic Acids Res 42, 8845–8860. 10.1093/nar/gku555.

16. Heumos, L., Schaar, A.C., Lance, C., Litinetskaya, A., Drost, F., Zappia, L., Lücken, M.D., Strobl, D.C., Henao, J., Curion, F., et al. (2023). Best practices for single-cell analysis across modalities. Nat Rev Genet 24, 550–572. 10.1038/s41576-023-00586-w.

17. Adil, A., Kumar, V., Jan, A.T., and Asger, M. (2021). Single-Cell Transcriptomics: Current Methods and Challenges in Data Acquisition and Analysis. Front Neurosci 15. 10.3389/fnins.2021.591122.

18. 18. Miwa, H., Dimatteo, R., de Rutte, J., Ghosh, R., and Di Carlo, D. (2022). Single-cell sorting based on secreted products for functionally defined cell therapies. Microsyst Nanoeng 8, 84. 10.1038/s41378-022-00422-x.

19. Holt, L.M., and Olsen, M.L. (2016). Novel Applications of Magnetic Cell Sorting to Analyze Cell- Type Specific Gene and Protein Expression in the Central Nervous System. PLoS One 11, e0150290. 10.1371/journal.pone.0150290.

20. Guo, W., Hu, Y., Qian, J., Zhu, L., Cheng, J., Liao, J., and Fan, X. (2023). Laser capture microdissection for biomedical research: towards high-throughput, multi-omics, and single-cell resolution. Journal of Genetics and Genomics 50, 641–651. 10.1016/j.jgg.2023.07.011.

21. Boulay, A.-C., Saubaméa, B., Declèves, X., and Cohen-Salmon, M. (2015). Purification of Mouse Brain Vessels. Journal of Visualized Experiments. 10.3791/53208.

22. Bernard-Patrzynski, F., Lécuyer, M.-A., Puscas, I., Boukhatem, I., Charabati, M., Bourbonnière, L., Ramassamy, C., Leclair, G., Prat, A., and Roullin, V.G. (2019). Isolation of endothelial cells, pericytes and astrocytes from mouse brain. PLoS One 14, e0226302. 10.1371/journal.pone.0226302.

23. Brzica, H., Abdullahi, W., Reilly, B.G., and Ronaldson, P.T. (2018). A Simple and Reproducible Method to Prepare Membrane Samples from Freshly Isolated Rat Brain Microvessels. Journal of Visualized Experiments. 10.3791/57698.

24. Paraiso, H.C., Wang, X., Kuo, P.-C., Furnas, D., Scofield, B.A., Chang, F.-L., Yen, J.-H., and Yu, I.-C. (2020). Isolation of Mouse Cerebral Microvasculature for Molecular and Single-Cell Analysis. Front Cell Neurosci 14. 10.3389/fncel.2020.00084.

25. Mi, J., Sun, A., Härtel, L., Dilling, C., Meybohm, P., and Burek, M. (2024). Isolation of Capillaries from Small Amounts of Mouse Brain Tissue. In, pp. 27–38. 10.1007/978-1-0716-3662-6_2.

26. Lee, Y.-K., Uchida, H., Smith, H., Ito, A., and Sanchez, T. (2019). The isolation and molecular characterization of cerebral microvessels. Nat Protoc 14, 3059–3081. 10.1038/s41596-019-0212-0.

27. Lau, K., Porschen, L.T., Richter, F., and Gericke, B. (2023). Microvascular blood-brain barrier alterations in isolated brain capillaries of mice over-expressing alpha-synuclein (Thy1-aSyn line 61). Neurobiol Dis 187, 106298. 10.1016/j.nbd.2023.106298.

28. He, L., Vanlandewijck, M., Raschperger, E., Andaloussi Mae, M., Jung, B., Lebouvier, T., Ando, K., Hofmann, J., Keller, A., and Betsholtz, C. (2016). Analysis of the brain mural cell transcriptome. Sci Rep 6, 35108. 10.1038/srep35108.

29. Vanlandewijck, M., He, L., Mae, M.A., Andrae, J., Ando, K., Del Gaudio, F., Nahar, K., Lebouvier, T., Lavina, B., Gouveia, L., et al. (2018). A molecular atlas of cell types and zonation in the brain vasculature. Nature 554, 475–480. 10.1038/nature25739.

30. He, L., Vanlandewijck, M., Mae, M.A., Andrae, J., Ando, K., Del Gaudio, F., Nahar, K., Lebouvier, T., Lavina, B., Gouveia, L., et al. (2018). Single-cell RNA sequencing of mouse brain and lung vascular and vessel-associated cell types. Sci Data 5, 180160. 10.1038/sdata.2018.160.

31. Adams, R.H., Wilkinson, G.A., Weiss, C., Diella, F., Gale, N.W., Deutsch, U., Risau, W., and Klein, R. (1999). Roles of ephrinB ligands and EphB receptors in cardiovascular development: demarcation of arterial/venous domains, vascular morphogenesis, and sprouting angiogenesis. Genes Dev 13, 295–306. 10.1101/gad.13.3.295.

32. You, L.-R., Lin, F.-J., Lee, C.T., DeMayo, F.J., Tsai, M.-J., and Tsai, S.Y. (2005). Suppression of Notch signalling by the COUP-TFII transcription factor regulates vein identity. Nature 435, 98–104. 10.1038/nature03511.

33. Cui, X., Lu, Y.W., Lee, V., Kim, D., Dorsey, T., Wang, Q., Lee, Y., Vincent, P., Schwarz, J., and Dai, G. (2015). Venous Endothelial Marker COUP-TFII Regulates the Distinct Pathologic Potentials of Adult Arteries and Veins. Sci Rep 5, 16193. 10.1038/srep16193.

34. Varga, C., Tamas, G., Barzo, P., Olah, S., and Somogyi, P. (2015). Molecular and Electrophysiological Characterization of GABAergic Interneurons Expressing the Transcription Factor COUP-TFII in the Adult Human Temporal Cortex. Cerebral Cortex 25, 4430–4449. 10.1093/cercor/bhv045.

35. Fuentealba, P., Klausberger, T., Karayannis, T., Suen, W.Y., Huck, J., Tomioka, R., Rockland, K., Capogna, M., Studer, M., Morales, M., et al. (2010). Expression of COUP-TFII Nuclear Receptor in Restricted GABAergic Neuronal Populations in the Adult Rat Hippocampus. The Journal of Neuroscience 30, 1595–1609. 10.1523/JNEUROSCI.4199-09.2010.

36. Tarhan, L., Bistline, J., Chang, J., Galloway, B., Hanna, E., and Weitz, E. (2023). Single Cell Portal: an interactive home for single-cell genomics data. bioRxiv, 2023.07.13.548886. 10.1101/2023.07.13.548886.

37. 37. Bjørnholm, K.D., Del Gaudio, F., Li, H., Li, W., Vazquez-Liebanas, E., Mäe, M.A., Lendahl, U., Betsholtz, C., Nilsson, P., Karlström, H., et al. (2023). A robust and efficient microvascular isolation method for multimodal characterization of the mouse brain vasculature. Cell Reports Methods 3, 100431. 10.1016/j.crmeth.2023.100431.

38. Yu, C., Young, S., Russo, V., Amsden, B.G., and Flynn, L.E. (2013). Techniques for the Isolation of High-Quality RNA from Cells Encapsulated in Chitosan Hydrogels. Tissue Eng Part C Methods 19, 829–838. 10.1089/ten.tec.2012.0693.

39. Autengruber, A., Gereke, M., Hansen, G., Hennig, C., and Bruder, D. (2012). Impact of enzymatic tissue disintegration on the level of surface molecule expression and immune cell function. Eur J Microbiol Immunol (Bp) 2, 112–120. 10.1556/EuJMI.2.2012.2.3.

40. Dempsey, M.E., Woodford-Berry, O., and Darling, E.M. (2021). Quantification of Antibody Persistence for Cell Surface Protein Labeling. Cell Mol Bioeng 14, 267–277. 10.1007/s12195-021-00670-3.

41. Li, S., Chen, L., Peng, X., Wang, C., Qin, B., Tan, D., Han, C., Yang, H., Ren, X., Liu, F., et al. (2018). Overview of the reporter genes and reporter mouse models. Animal Model Exp Med 1, 29–35. 10.1002/ame2.12008.

42. Yamazaki, Y., and Kanekiyo, T. (2017). Blood-Brain Barrier Dysfunction and the Pathogenesis of Alzheimer’s Disease. Int J Mol Sci 18. 10.3390/ijms18091965.

