## Supplementary Figure for "Optimized Enrichment of Murine Blood-Brain Barrier Vessels with a Critical Focus on Network Hierarchy in Post-Collection Analysis"

**SUPPLEMENTAL INFORMATION**

**Supplemental Figures 1 and 2.** Original western blots in support of Figure 3.

**Supplemental Figure 3. Calponin1 (*Cnn1*) Expression Analysis of Output Material from each Protocol Underscores the Utility of Filtration in Reducing Non-Capillary Vessels.** *Cnn1* transcripts were evaluated using qRT-PCR. *Tbp* was our housekeeping gene, and we normalized the measurements for each protocol group separately to better appreciate the fold difference in transcript levels between them. Specifically, enriched samples (Group 2) were referenced to whole cortex samples (Group 1) (left graph), and enriched and filtered samples (Group 3) were referenced to enriched samples (Group 2) (right graph). Dotted lines indicate “1” for the reference sample. Individual data points are shown with averages represented by bars – Whole Cortex (Group 1): yellow bars, Enriched by DGC (Group 2): blue bars, and Enriched and Filtered (Group 3): pink bars. Error bars are standard deviation, and * denotes $p \leq 0.05$ by Student’s T-test.

**Supplemental Figure 1.**

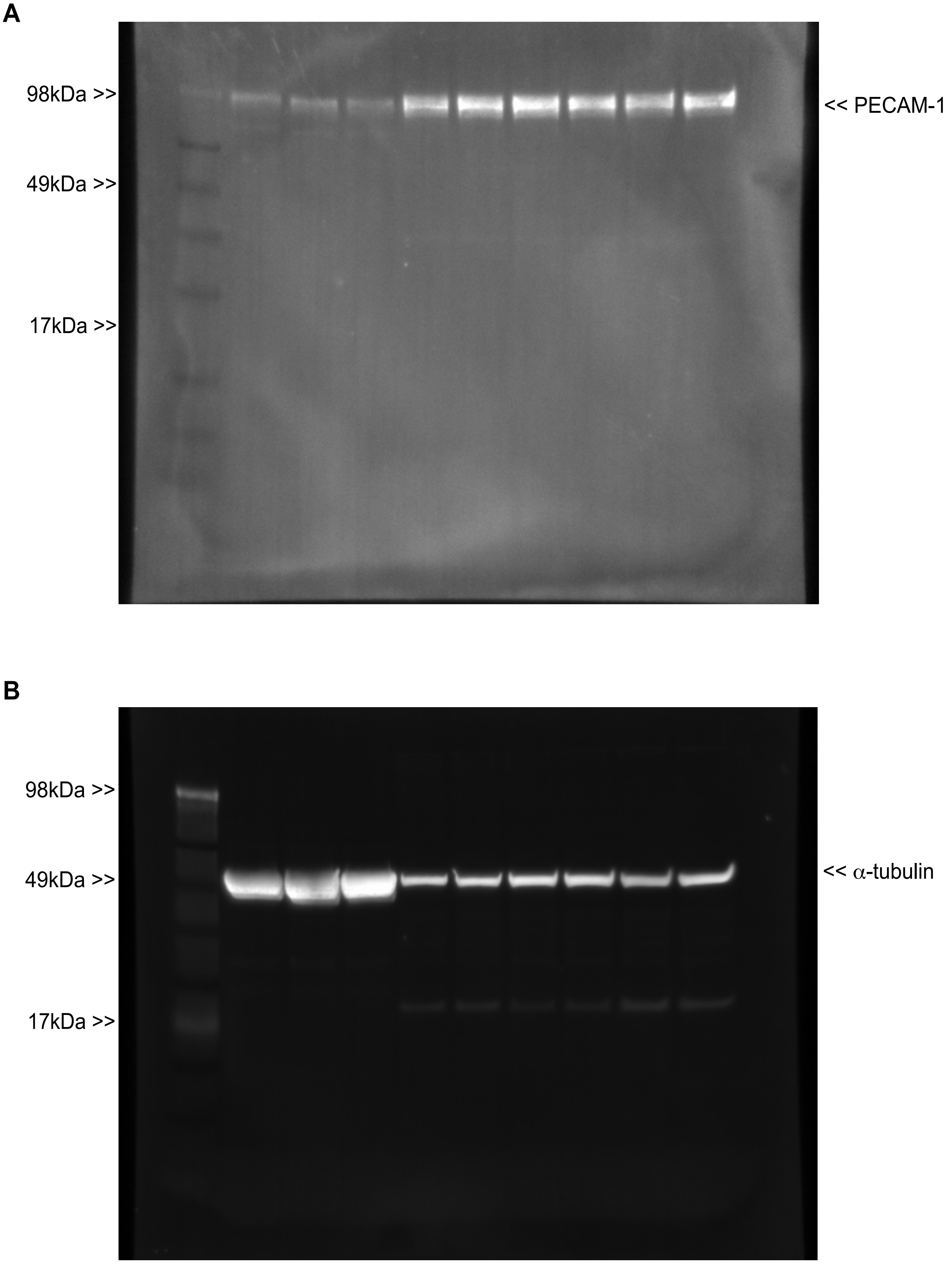

**Supplemental Figure 2.**

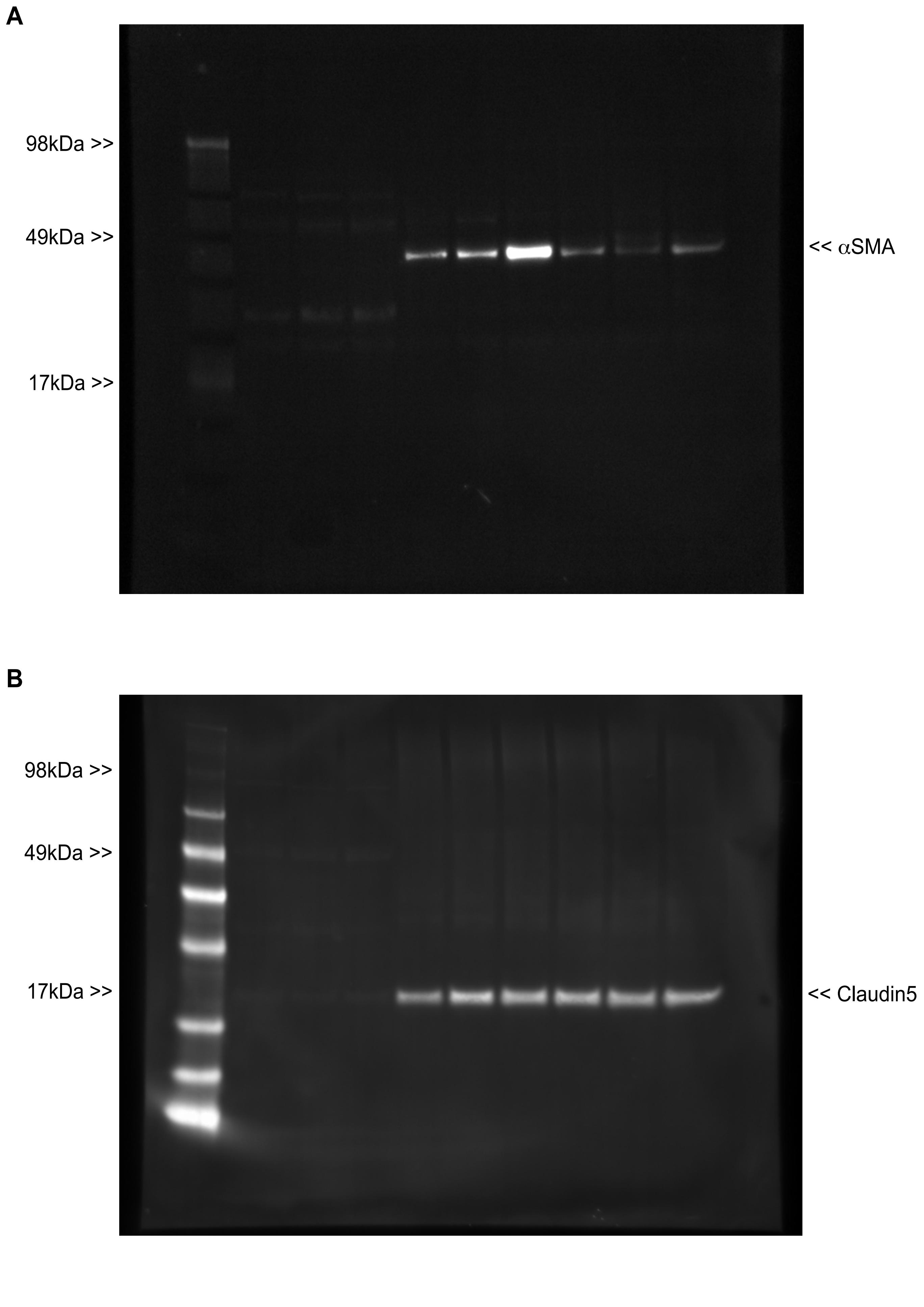

### Supplemental Figure 3.

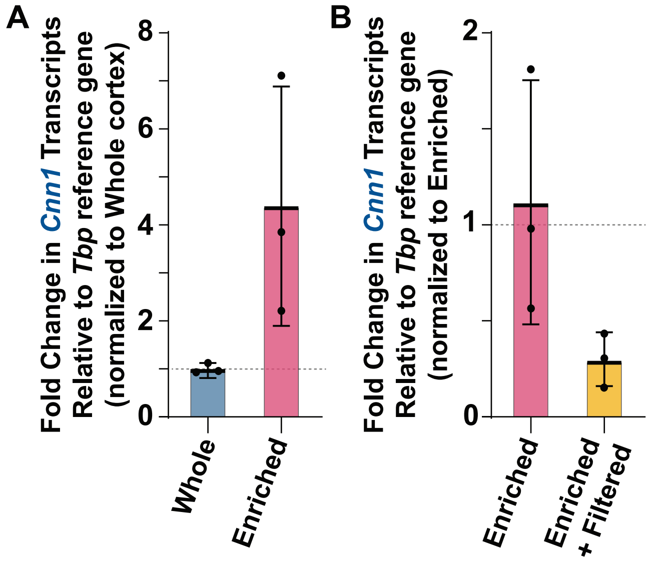
